# A nitrogen assimilation bottleneck can limit *Bacillus subtilis* growth in plant culture media

**DOI:** 10.64898/2026.03.30.715200

**Authors:** Avaniek Cabales, Rachel Warthen, Harsh Bais, Aditya M. Kunjapur

**Affiliations:** Department of Chemical and Biomolecular Engineering, University of Delaware, Newark, DE, USA; Department of Plant and Soil Sciences, University of Delaware, Newark, DE, USA

**Keywords:** Bacillus subtilis, surfactin, nitrogen assimilation, plant culture media, hydroponic cultivation, rhizobacteria, plant growth promotion, amino acids

## Abstract

Microbial engineering offers potential for improving the sustainability of agriculture by providing greater control of desired microbial functions. However, successful control of engineered functions requires greater understanding of their robustness under diverse conditions including those used for plant hydroponics. Here, we studied biomass accumulation and surfactin biosynthesis by an engineered derivative of *Bacillus subtilis* PY79 in common plant culture media as a model system for interrogating metabolic robustness. We report the observation that PY79 and all other *B. subtilis* strains that we tested, including natural isolates, exhibited difficulty growing under shaking incubation in defined media where the only nitrogen sources were inorganic. In contrast, assimilation of inorganic nitrogen sources functioned relatively robustly under static incubation in these same media. Our findings may offer some guidance for use of *B. subtilis* in controlled environment agriculture and could aid future efforts to identify the molecular basis for the agitation-dependent effect on nitrogen assimilation.

## Introduction

While the biotechnology industry has historically exploited microbial engineering for use in bioreactors, advances in microbial engineering have demonstrated an increasing number of compelling proof-of-concepts in less well-controlled or uncontrolled environments such as in soil or in the rhizosphere (1, 2). In many of these environments, microbial species naturally contribute substantially to the health of their plant host (3, 4). In principle, engineering could augment the abilities of natural microbes by expanding the breadth of functions that they could perform or by achieving desired regulation of their functions across space or time (1). However, to successfully implement microbial engineering for these purposes, we must gain greater understanding of the context-dependence of microbes in these environments. This understanding begins with basic nutrient assimilation pathways and the regulation of these pathways to achieve desired engraftment or exclusion of the engineered microbe based on the availability or lack of an available metabolic niche (5, 6).

Modern agriculture relies heavily on pesticides and fertilizers, negatively impacting the sustainability of global food production (4). While plant growth promoting rhizobacteria (PGPR) have the promise to serve as a more sustainable alternative, natural isolates used to enrich foreign soils often yield inconsistent results (7). The rhizosphere is an uncontrolled and complex environment much different from laboratory testing conditions, with variability stemming from numerous factors such as plant variety, environmental conditions, and soil composition (3). To study plant-microbe interactions in a more controlled environment with high throughput and reproducibility, one can consider the use of plant culturing in liquid media under a gnotobiotic setup. The insights gained from studying plant-microbe interactions in plant culturing media may also translate to controlled environment agriculture, where hydroponic systems are common.

Beneficial microbes including *Bacillus subtilis* have been added to hydroponic systems and are of rising interest given their promise of improving nutrient assimilation for plant hosts and limiting microbial pathogens (8–12).

As natural soil bacteria, wild-type *B. subtilis* strains form biofilms in response to plant polysaccharides or secretion of simple of carboxylic acids, giving them an ability to stably colonize plant roots (13, 14). Many strains of *B. subtilis* produce diverse secreted small molecules that promote plant growth and suppress disease, including cyclic-lipopeptides such as surfactin (15). *B. subtilis* is also poised well for further translation in agricultural biotechnology given its genetic tractability, history in industrial fermentation (16), and formation of endospores that exhibit high stress tolerance and enable shelf-stable formulations for seed coating (17) or foliar spraying (18). Similarly, many *B. subtilis* strains have been designated as benign by the US EPA and Generally Recognized as Safe (GRAS) by the FDA (19, 20). These factors likely contribute to the active licensing of *B. subtilis* isolates such as strain UD1022 by large agrochemical companies (21).

Surfactin is a model metabolic target for engineering controllable overproduction due to its many plant benefits, which include improving colonization of beneficial bacteria (22–24), exerting antimicrobial effects on plant pests (25–27), inducing systemic resistance (28), and increasing drought tolerance (29). While many domesticated strains of *B. subtilis* have lost their capacity to produce surfactin, the phenotype can be restored by introduction of a full-length *sfp* gene, which encodes a phosphopantetheinyl transferase required for activation of surfactin synthetases (30). Surfactin overproduction has been achieved in bioreactors through genetic modifications that increase expression of the *srf* operon and flux to surfactin precursors, including branched chain amino acids and fatty acids (23, 31–33). Surfactin biosynthesis has also been improved by altering fermentation conditions (32, 34–36), including careful control of carbon and nitrogen sources under high density fermentation conditions (37). The role of surfactin in plant growth, colonization, and plant growth promotion is still under debate (38–40), partly due to the strong dependence of *in situ* surfactin biosynthesis on plant co-culture conditions. Furthermore, surfactin titers show high variability across *Bacillus subtilis* strains (41). In general, it could be the case that a different balance of surfactin biosynthesis and host fitness would be optimal in a bioreactor than in hydroponic culture or the rhizosphere as an effective bioinoculant would need to compete among a microbiome and associate with the plant root. Additionally, it is also likely that lower titers than those achieved in large-scale fermentations would be sufficient for benefiting plant growth if produced locally on the root surface (40). Thus, surfactin represents a simple and model chemistry for study given that *B. subtilis* PY79 offers a surfactin negative blank canvas and that ample knowledge is available to tune surfactin productivity.

Towards the goal of understanding how *B. subtilis* PY79 and biosynthesized surfactin could interact with plants, and how *B. subtilis* strains might grow in plant culture media conditions, we sought to culture engineered strains under conditions that increasingly mimicked plant hydroponic culturing conditions, in the absence of a plant. We were surprised to observe differential growth patterns in an agitation-dependent manner that were conserved among all tested strains. Our study highlights the importance of organic nitrogen sources for robust growth of *B. subtilis* in common plant culturing media under conditions that promote greater aeration.

## Results

### Growth and surfactin biosynthesis of engineered PY79 sfp^+^ using common bacterial growth media

Before we began to engineer PY79 to produce surfactin, we sought to understand the potential toxicity of our desired product on the microbial host, independent of the potential metabolic burden associated with surfactin biosynthesis. Thus, we began by investigating how the concentration of exogenously supplemented surfactin would affect the fitness of PY79. We cultured the progenitor PY79 in either LB or S750 media at concentrations up to 0.8 g/L and observed no differences among strain growth in either media condition (**Fig. S1**). Most of these concentrations are well above what is necessary for induction of plant benefits, with as little as 0.025 g/L exogenous surfactin demonstrating growth benefits to tomato plants, for example(42).

As briefly mentioned earlier, PY79 is a surfactin null strain due to a non-functional *sfp* gene(43). We restored surfactin biosynthesis by genome integration of the *sfp* gene under constitutive control by the P_veg_ promoter into the *lacA* locus (**Fig. 1a**). Importantly, this PY79 sfp^+^ strain relies on the native *srf* operon and its regulation for the assembly of surfactin (**Fig. 1b**). After constructing this strain, we investigated its ability to biosynthesize surfactin under various conditions, which we could then measure by HPLC-UV, where we observed three distinct peaks in our commercial standard. These peaks were present in the supernatant of PY79 sfp^+^, but not in the supernatant of PY79 (**Fig. 1c**).

**Figure 1.**
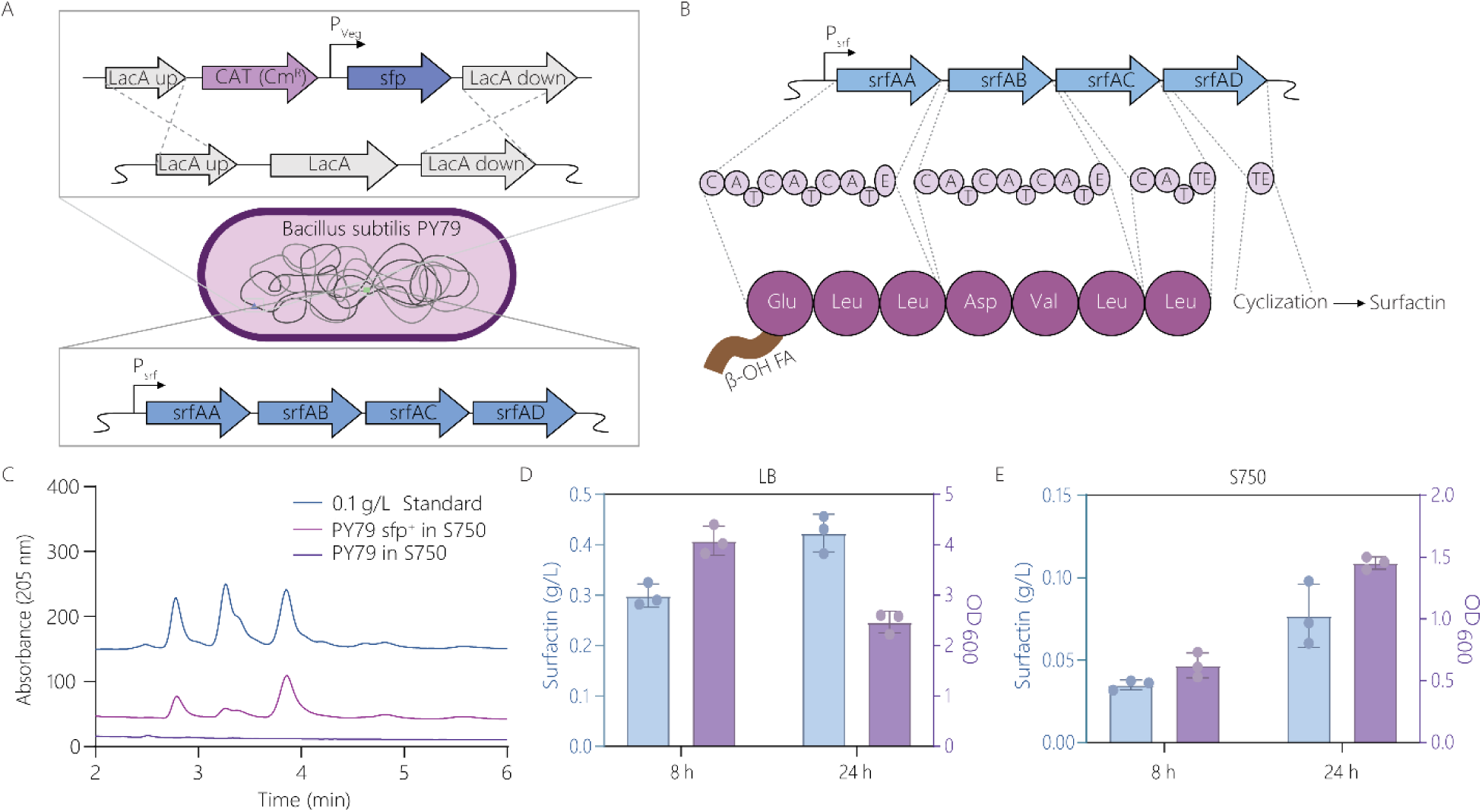
Growth and surfactin biosynthesis of engineered PY79 sfp^+^ using common bacterial growth media. (a) Surfactin positive (PY79 sfp^+^) strain was constructed by introducing *sfp* under vegetative control in the LacA locus. The protein encoded by this gene activates the other proteins in the srfA operon through post-translational modification. (b) Proteins encoded in the *srfA* operon linearly append amino acids starting with a fatty acid until SrfAD forms the lactone ring. (c) Surfactin chromatograms of the commercial standard, supernatant of sfp^+^ grown in S750, and PY79 in S750. Growth of sfp^+^ strain and surfactin production in (d) rich LB media and (e) minimal S750 media.

To begin to understand the context-dependence of the growth of our strain and its surfactin productivity, the first variable we studied was the influence of the growth media during shaking incubation at 37°C that is typical for batch cultivation of bacterial monocultures under laboratory conditions. When we cultivated the PY79 sfp^+^ strain in LB media, we saw an average of 0.42 g/L surfactin produced by an endpoint of 24 h and a final OD_600_ above 2.0 (**Fig. 1d**). In contrast, when we cultivated the PY79 sfp^+^ strain in S750 minimal media, which contains 20 mM inorganic nitrogen from ammonium, 10 mM of glutamate, and 1% glucose as a carbon source, we observed decreased biomass accumulation and much lower surfactin production. While the final OD_600_ was nearly 1.5, we observed only an average of 0.077 g/L surfactin produced by 24 h, indicating that the productivity of surfactin biosynthesis normalized to the level of biomass formation had declined substantially in the minimal media condition under shaking conditions. (**Fig. 1e**).

We also compared production under these conditions for the wild-type strain UD1022, which is a sfp^+^ strain that is thought to naturally produce surfactin. However, we were unable to detect surfactin in LB and S750 media under conditions tested, despite growth of up to optical densities of 3.5 and 1.3 in LB and S750 respectively (**Fig. S2**). Based on these results, we proceeded to evaluate the context dependence of growth and surfactin production by focusing on just the engineered PY79 sfp^+^ strain and by advancing towards plant culturing protocols.

### L-glutamate supplementation substantially improves growth under common plant culture media

After demonstrating surfactin production under typical laboratory conditions, we moved to media conditions that we deemed more representative of plant culturing in laboratory environments, namely the common plant culture media half-strength Murashige and Skoog (½X MS) supplemented with 1.5% sucrose. Initially, we looked at growth of PY79 in this media and maintained a temperature of 37°C and provided shaking incubation to promote microbial growth in the absence of a plant. However, we noted poor growth in ½X MS media with 1.5% sucrose (**Fig. 2a**). Although ½X MS media contains 825 mg/L of ammonium nitrate and 950 mg/L potassium nitrate (as well as 1 mg/L glycine), we wondered whether PY79 was facing a limitation in carbon or nitrogen availability. We noted that S750 minimal media contains 1.32 g/L ammonium sulfate as well as 10 mM L-glutamate because past personal communications suggested that the addition of L-glutamate would improve the robustness and reproducibility of growth. L-glutamate has potentially multifaceted contributions to the growth of *B. subtilis* as it has a well-established role in nitrogen metabolism (44, 45), is incorporated into the secreted poly-γ-glutamic acid (PGA) polymers that contribute to biofilm formation (46, 47), and is a component of surfactin and could be serving as a carbon source. Additionally, in the natural context of root colonization, root exudates often contain primary metabolites including amino acids which serve as chemoattractants driving root colonization patterns (48–50), and recently a prominent role for the related amino acid L-glutamine was reported as an influential driver of root microbial colonization upon local endodermal leakage(51). For these reasons, we tested the influence of L-glutamate supplementation on biomass accumulation in the ½X MS medium. We also tested the influence of sucrose addition or combined L-glutamate and sucrose addition on biomass accumulation to better determine whether nitrogen or carbon was growth-limiting. Our results clearly indicate that carbon was not limiting because 10 mM L-glutamate supplementation substantially improved strain growth whereas 15 g/L sucrose addition did not **(Fig. 2a)**.

**Figure 2.**
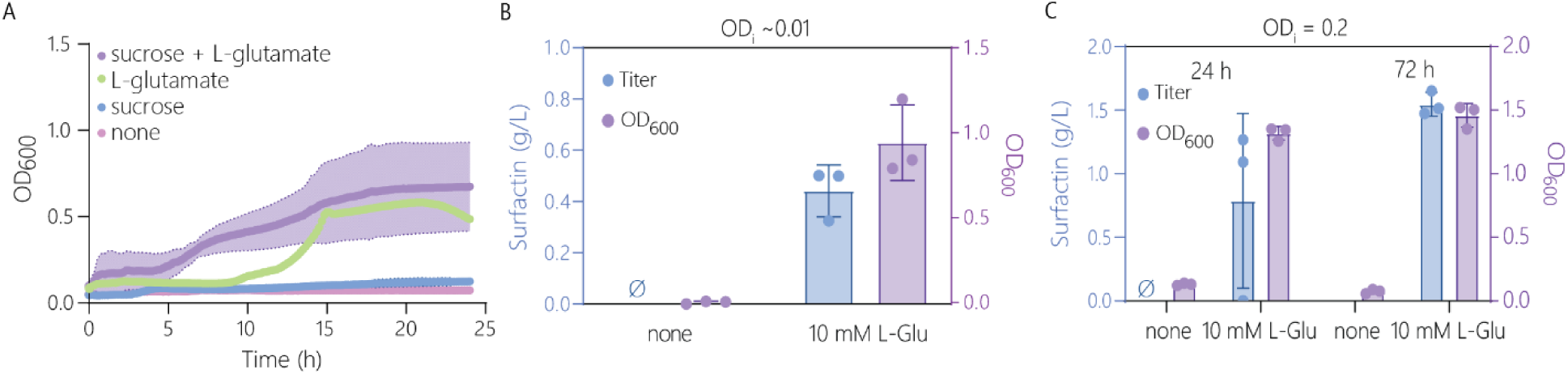
L-glutamate supplementation substantially improves growth under common plant culture media. (a) Growth of PY79 in ½X MS media supplemented with 10mM glutamate and/or 1.5% sucrose over 24 hours. (b) Growth and surfactin production of PY79 sfp^+^ strain over 24 hours at a low inoculation ratio. (c) Growth and surfactin production at 24 and 72 hours of sfp^+^ strain after inoculation at a higher OD of 0.2. Null sign indicates no detectable quantities were observed.

Similarly, under shaking conditions at 37°C, we observed minimal growth of PY79 sfp^+^ and detected no surfactin production without glutamate addition (**Fig. 2b**). Having established an important role for L-glutamate supplementation under these conditions, we proceeded to test its influence on final culture OD and surfactin titer, including when using two distinct ODs of inoculation. When we supplemented 10 mM L-glutamate under our initial experimental conditions inoculating at a ratio of 1:100 from cells grown to confluence, we observed surfactin production at the rate of 0.44 g/L at 24 h (**Fig. 2b**). Given that surfactin production is cell density-dependent (52), we also tested inoculation at an initial OD of 0.2 and incubated for 72 h. We observed higher OD (nearly 1.5) and much higher surfactin titers of 1.5 g/L (**Fig. 2c**), which are 3.6-fold higher than the surfactin titers that we obtained when using LB medium. This suggests that the reason we observed low surfactin titers in the previously described minimal media experiments performed for 24 h may have been because of insufficient cultivation under those conditions.

### Characterizing the L-glutamate-dependent effect by varying concentration and amino acid identity

We next sought to understand the concentration-dependent influence of L-glutamate on surfactin biosynthesis as well as whether other amino acids could provide similar benefit. For these experiments, we observed that surfactin titers did not change appreciably between 24 h and 72 h (**Fig. 3a**). Here, we found that L-glutamate supplementation in the range of 1-10 mM results in comparable surfactin titers of nearly approximately 0.8 g/L and OD of approximately 1. At 0.5 mM, we observe a decrease in final OD_600_ to 0.49 and surfactin production of 0.15 g/L. At 72 hours, the trend remains similar with slightly more surfactin production for cases with 1 mM or greater L-glutamate addition **(Fig. 3b)**. This is consistent with our earlier observation suggesting that nitrogen assimilation is limiting growth.

**Figure 3.**
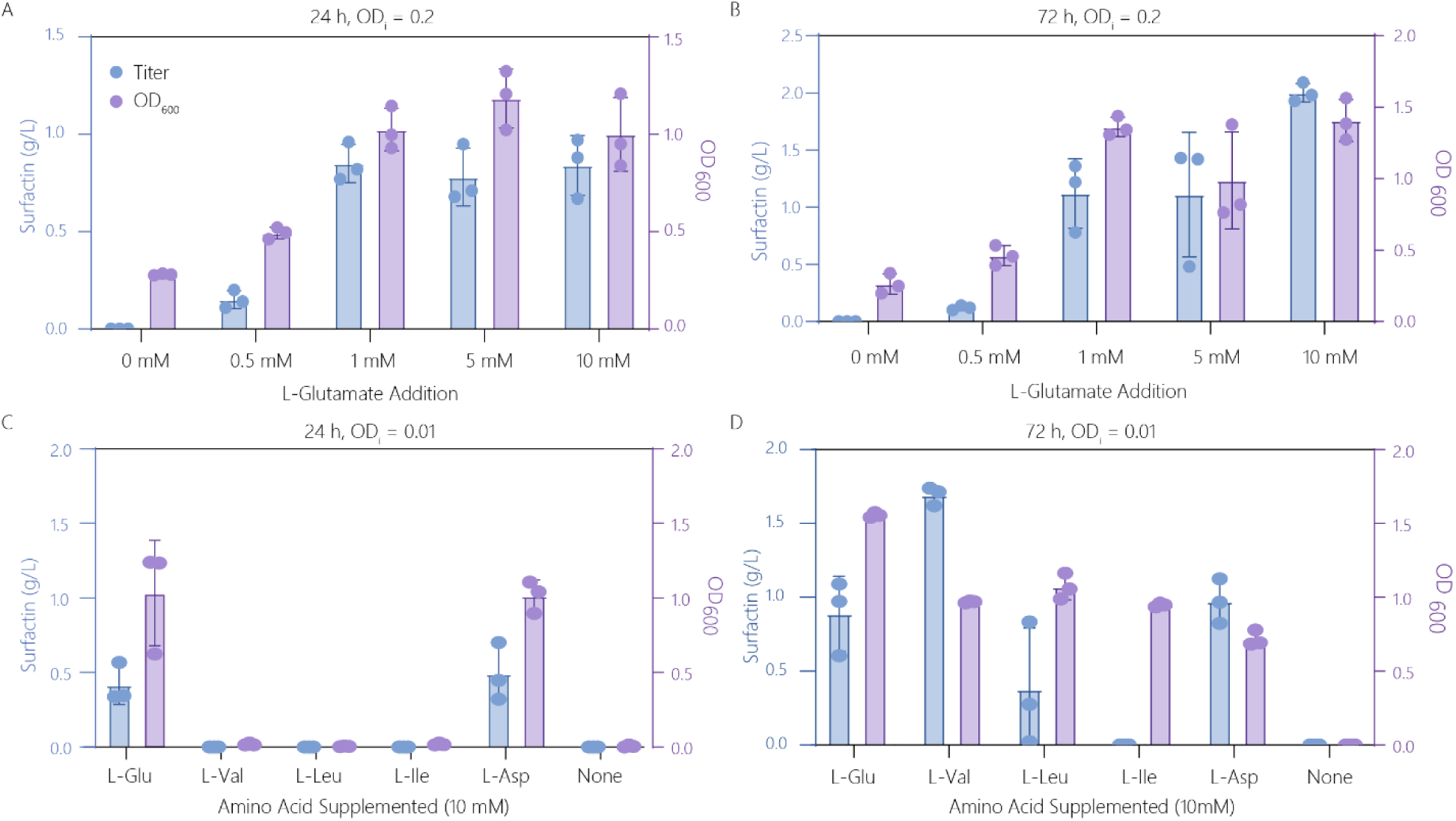
Characterizing the L-glutamate-dependent effect by varying concentration and amino acid identity. Growth and surfactin production of sfp^+^ strain with varying supplemented glutamate over (a) 24 h and (b) 72 h. Results of varying amino acid supplemented over (c) 24 h and (d) 72 h.

To next disentangle whether L-glutamate is serving a unique role or a more general role as an organic nitrogen source that could be replaced by other amino acids, we supplemented 10 mM of L-aspartate, L-isoleucine, L-leucine, or L-valine. We chose these amino acids as they compose the ring structure in surfactin and could thereby contribute to final OD and surfactin titer. We tested inoculation OD of either 0.01 or 0.2 and measured growth and surfactin production over 72 h (**Fig. 3c-d**). At lower inoculation ratios, L-glutamate and L-aspartate were able to grow to an OD of approximately 1.0 and produced nearly 0.5 g/L of surfactin within 24 h; however, by 72 h, any of these amino acids were able to support the growth of the strains. Similarly, except for L-isoleucine, the amino acids that we tested resulted in similar final titers of surfactin. We observed similar trends when using the larger inoculation OD, except in the case of isoleucine, which resulted in inconsistent surfactin titers (**Fig. S3).**

### Comparing shaking or static incubation and nitrogen assimilation across different *B. subtilis* strains

Our experiments thus far highlighted the difficulty of *B. subtilis* PY79 sfp^+^ to assimilate inorganic nitrogen sources in the context of the plant hydroponic media. Next, we sought to consider the effects of lower temperatures and static incubation. Lower temperature (25°C) is more realistic for hydroponic plant growth, whereas we considered static incubation possibly less representative for hydroponic culturing but still interesting given that in industry plants may be kept static while liquid media would likely flow, resulting in a potentially wide range of agitation conditions experienced by bioinoculants. Going into these experiments, we hypothesized that lower temperature and lower agitation rates would each be likely to slow microbial biomass accumulation and result in decreased surfactin production.

First, we cultivated our engineered strain at lower temperatures of 25 °C under conditions of shaking incubation (1000 rpm) to isolate the effects of lower temperature. Under these conditions, growth in the culture remained dependent on inclusion of an organic nitrogen source **(Fig. 4a-b).** Without supplemental L-glutamate, we detected no surfactin production and poor growth at these lower temperatures; however, when supplementing glutamate at 10 mM of L-glutamate, we detect about 0.05 g/L surfactin and growth to approximately 1.0 OD, regardless of initial inoculation OD. With addition of only 1 mM L-glutamate, we see reduced growth with final ODs of 0.35 or 0.16 at higher and lower inoculations densities, respectively. We observed much poorer surfactin production, seeing approximately 0.055 g/L at high inoculation densities and 10 mM L-glutamate at the lower temperatures as compared to our results at 37 °C.

**Figure 4.**
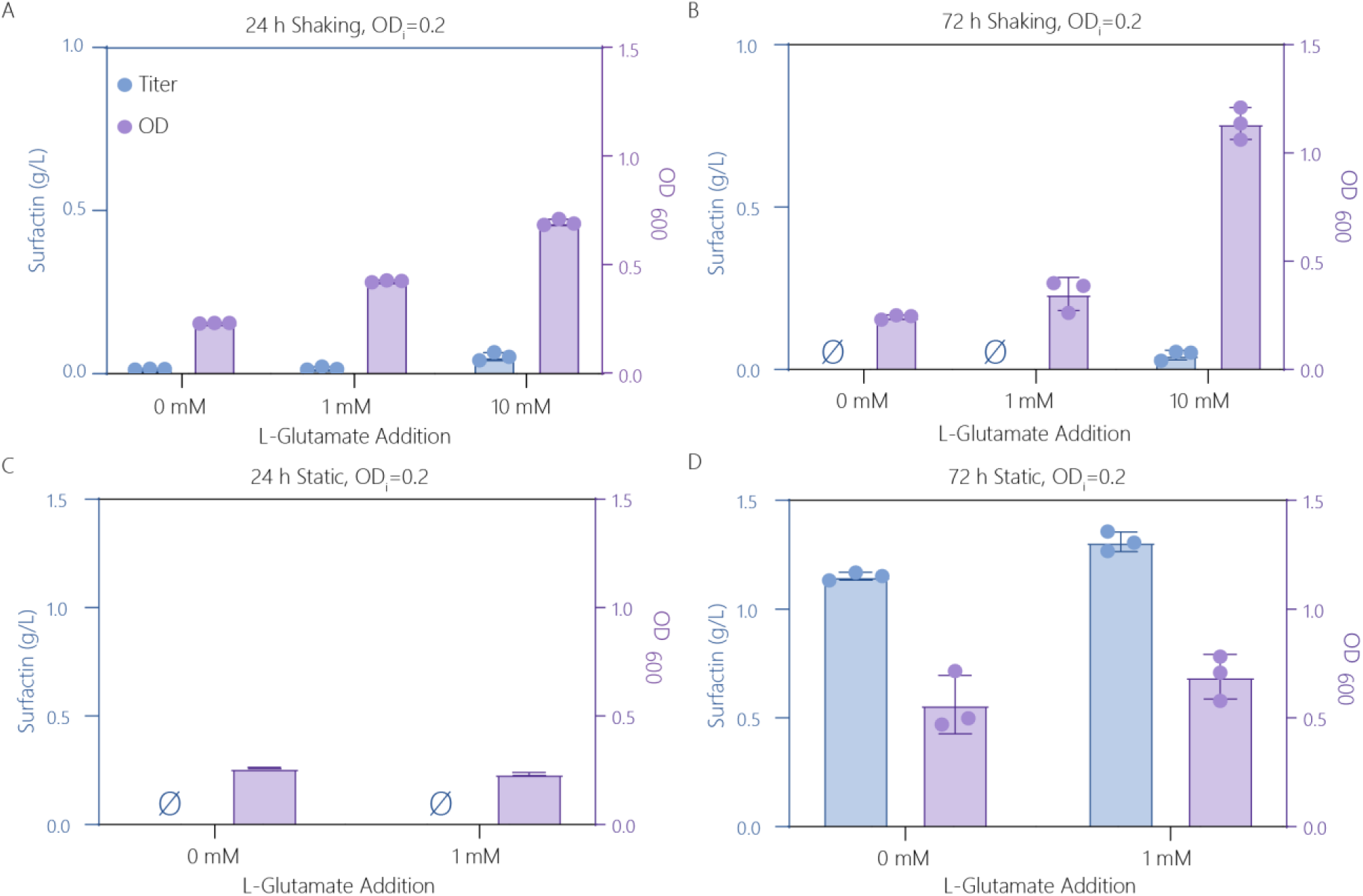
The importance of organic nitrogen provision on *B. subtilis* PY79 sfp^+^ biomass accumulation varies across shaking and static incubation. Growth and surfactin production of sfp^+^ strain under shaking conditions at 25 °C over (a) 24 h and (b) 72 h, and under static conditions at (c) 24 h and (d) 72 h. Null sign indicates no detectable quantities were observed.

In moving towards conditions more relevant to hydroponic conditions, we then tested under static conditions at 25 °C, with and without supplementation of 1 mM glutamate. To our surprise, we observed robust growth and production of surfactin **(Fig 4c-d)**. Within 24 h, culture OD reaches 0.26 and 0.23, but with no detectable surfactin produced. However, when we look to 72 h timepoints while supplementing no or 1 mM L-glutamate, we see growth and production with final ODs of approximately 0.6 and concentrations of 1.2 and 1.3 g/L. Thus, under static conditions, regardless of the addition of organic nitrogen, we noted growth and surfactin production by 72 hours. Collectively, these observations suggest that the assimilation of inorganic nitrogen by *B. subtilis* could be differentially regulated based on abiotic factors that change during agitation, such as dissolved oxygen concentration or possibly mechanical force.

To better understand the potential broader applicability and impact of this observation for other *B. subtilis* strains rather than just the PY79 sfp^+^ strain, we next expanded our investigation to include the laboratory *B. subtilis* strains 168 and W23 as well as a plant-growth promoting isolate UD1022(21). We chose 168 and W23 due to their phylogenetic relationship to PY79, where PY79 resulted from a cross with 168 and W23 resulting in a strain similar to 168 with W23 islands(53). To gain a more comprehensive understanding, we tested these strains with a variety of organic and inorganic nitrogen sources, in the presence and absence of agitation. *B. subtilis* 168 is an L-tryptophan auxotroph, so we supplemented 50 µg/mL L-tryptophan to the media only for 168 samples. We included an L-tryptophan control for the other strains to determine whether this additional nitrogen would be enough organic nitrogen to support the growth of the strains. Using a base 1/2X MS with 1.5% sucrose media, we supplemented an additional 10 mM of either an organic or inorganic nitrogen source (**Fig. 5a-b**).

**Figure 5.**
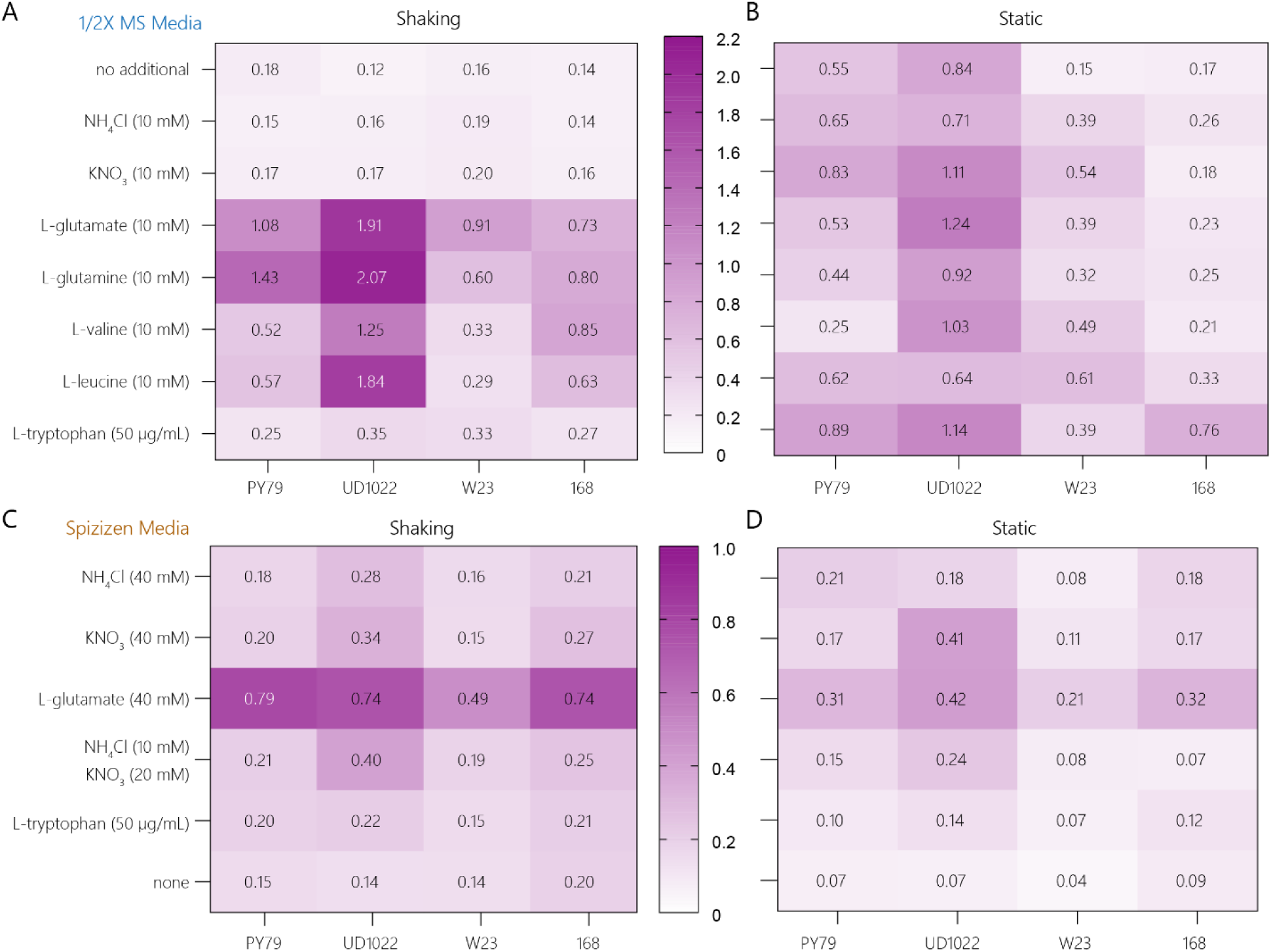
The observed dependence on organic nitrogen assimilation under shaking incubation extends across all tested *B. subtilis* strains and across media. Growth of *B. subtilis* strains PY79, UD1022, W23, and 168 at 25 C in 1/2X MS media supplemented with varying nitrogen sources under (a) shaking or (b) static incubation. Growth of these strains in Spizizen minimal media while varying nitrogen source and incubating under (c) shaking or (d) static conditions. In Spizizen media, the strains all exhibited difficulty growing under static incubation even with L-glutamate provided.

We observed that both PY79 (sfp^-^) and UD1022, under static incubation, especially at 72 h, grow with the supplemented inorganic nitrogen or just the base 1/2X MS with 1.5% sucrose media. However, these strains required supplementation of the organic nitrogen sources to grow under shaking incubation. For 168, we saw the same trend under the shaking incubation. However, under static incubation, we only observed growth when supplementing cultures with an additional 50 µg/mL (resulting in 100 µg/mL total L-tryptophan supplied). This suggests that 50 µg/mL was a limiting concentration of L-tryptophan for 168 under static incubation but not under shaking incubation. For W23, we see the same general trend as PY79 (sfp^−^) and UD1022, but it did not grow to as high an OD, especially as compared to UD1022. We expect that UD1022 outgrows both PY79 and W23, especially under static contexts, due to lack of domestication and robust biofilm formation ability(54, 55). Collectively, our experiments revealed that, under the conditions we tested, there is indeed a broad tendency of *B. subtilis* strains to struggle with assimilation of inorganic nitrogen, but only under shaking incubation.

Finally, to determine if this phenomenon could be specific to our choice of 1/2X MS media, we tested these strains in Spizizen minimal base media. Spizizen media is commonly used for culturing *B. subtilis* under lab conditions. We tested 40 mM of ammonia, nitrate, or L-glutamate as sole nitrogen sources. Additionally, we included a condition with 10 mM ammonia and 20 mM nitrate, which is equivalent to the nitrogen in base 1/2X MS media. For each condition, we maintained sucrose as the carbon source, supplemented at 1.5% **(Fig. 5c-d)**. We noted that the strains grew poorly on inorganic nitrogen sources under shaking conditions, with growth to approximately 0.8 OD with supplemented glutamate for PY79, UD1022, 168. Similarly, W23 grew the best in this condition to OD 0.49. However, when inorganic nitrogen sources were supplied, PY79, W23, and 168 reached a final OD of approximately 0.2. UD1022 reached a max final OD of 0.4 on the mix of ammonia and nitrate but had a strong growth preference on the organic nitrogen. However, in this case, static conditions did not seem to improve growth on inorganic nitrogen, reaching final ODs less than 0.2 for PY79 and 168. W23 grew poorly in all cases of static incubation. To summarize, all strains appeared to struggle to assimilate inorganic nitrogen under all tested conditions when provided the Spizizen minimal base media. In contrast, in 1/2X MS media, we observed growth on inorganic nitrogen under static conditions.

### Analysis of published transcriptomic data pertaining to nitrogen assimilation

To ascertain how the differences between static and shaking incubation could manifest in differential expression of genes known to mediate inorganic nitrogen assimilation, we analyzed previously published transcriptomics results of a prototrophic lab strain of *B. subtilis* cultivated under a wide variety of conditions (56). Specifically, we wondered whether downregulation of genes required for inorganic nitrogen assimilation, as either nitrate or ammonia, under shaking incubation could be responsible for our observations. We compared the relative transcript levels of a variety of genes related to nitrogen metabolism, transport, and regulation in static or shaking incubations (56, 57) (**Fig. S4a-b)**. These cultures were grown in MSgg media, a minimal media containing 0.5% glycerol, 34 mM L-glutamate. We directly compared changes in expression levels in shaking for 24 h versus static for 36 h in MSgg media **(Fig. S4c)**. In comparing static cultures relative to the shaking condition, we noted *amtB*, *nasA*, and *glnK* were the most downregulated and that *gltA* and *gltB* were upregulated. As part of a two-component system with GlnL, GlnK regulates *gltT* for glutamate and aspartate uptake and *glsA* for glutamine degradation. AmtB and NasA transport ammonia and nitrate, respectively. In particular, the downregulation of *gltA* and *gltB* under shaking conditions could potentially contribute to the difficulty of cells to grow under these conditions. We briefly attempted to test this hypothesis by placing *gltAB* under inducible control; however, we were unable to obtain a viable colony, which could mean that the non-native control was detrimental to the fitness of the strain, though there are many other plausible explanations.

## Discussion and Conclusion

Microbial engineering holds the potential to engineer improved plant growth promotion in rhizobacteria. Evaluating the context dependence will be critical for utilizing biotechnology in environments that are very different than laboratory environments and highly dynamic. The rhizosphere is a highly complex and dynamic environment, whereas hydroponic culturing systems represent a more well-controlled environment and are of rising interest for the study of plant-microbe interactions and for industrial controlled-environment agriculture such as vertical farming. In such contexts, the use of bioinoculants is in its infancy yet gaining traction for purposes such as greater nutrient assimilation, limiting salinity stress, and disease control (58, 59). Towards this, we measured both the growth and desired function of a minimally engineered strain of *B. subtilis*, using biosynthesis of surfactin as a model compound for understanding plant-benign microbe interaction. During this process, due to stark differences in observed growth of *B. subtilis*, our attention shifted to the importance of organic versus inorganic nitrogen availability and utilization.

This work uncovered findings that could be surprising to many in the community as they are not well documented in the literature. The most important of these was that *B. subtilis* strains struggle to assimilate inorganic nitrogen under shaking incubation and are better able to assimilate it during static incubation in ½X MS media. To our knowledge, only one recent study has explicitly documented that *B. subtilis* exhibits difficulty assimilating inorganic nitrogen sources such as NH_4_Cl, which was observed with *B. subtilis* strain 168 under aerobic conditions in a 5 L bioreactor (60). Our work corroborates this observation under a different context, discovers that the ability to assimilate inorganic nitrogen sources depends upon agitation level, and extends this finding broadly across other domesticated and undomesticated strains of *B. subtilis*. In addition, we compared the assimilation of nitrogen to strain ability to biosynthesize surfactin under static and shaking conditions.

Nitrogen regulation is a complex balance of utilizing resources and adapting to changing nutrient availability, all while governing a critical process for microbial survival. Despite this, we observed poor adaptation of all tested *B. subtilis* strains to inorganic nitrogen when cultivated under agitation in the media and other cultivation conditions that we tested. We note that the conditions of the rhizosphere are starkly different than what we tested in numerous ways, but that some roles that plants serve with respect to nitrogen availability are interesting to consider. Within the rhizosphere, the form of nitrogen available can significantly affect microbial composition and behaviors (61, 62); for example, L-glutamine serves as an important signal for root colonization (63). In the rhizosphere, plants would secrete chemoattractants and metabolites to promote root colonization (64). However, hydroponic systems involve more agitation resulting from the flow of the growth medium in the system. In our experiments, we noted a difference in biomass in comparing agitated versus non-agitated conditions in ½X MS media with 1.5% sucrose, which may have implications for microbe colonization and growth behavior in these hydroponic systems as a function of the flow rate and agitation within the system. In a recent study that we conducted in parallel, we also observed that exogenous supplementation of L-glutamate and surfactin affected root colonization patterns in tomato seedlings (65).

Our study lays important groundwork for future studies to determine the molecular basis for this regulation. We performed differential gene expression analysis to explore the contributions of agitation to genes responsible for nitrogen assimilation, and we observed downregulation of *gltAB,* which assimilates ammonia into glutamate. Besides this, agitation could be perturbing nitrogen assimilation in other important ways. Since ½X MS media is uncommon for *B. subtilis* culturing and contains high metal concentrations that could increase formation of reactive oxygen species (ROS) when oxygen availability increases, we also tested Spizizen media, which has a long history of use for *B. subtilis* culturing and lower metal concentrations, though organic nitrogen sources in various forms, such as casein hydrolysate, are classically supplemented to Spizizen media, even when investigating nitrogen limitation (66–69). Because we observed limited inorganic nitrogen assimilation also when using Spizizen media, and because the default media includes casein hydrolysate supplementation, it is less likely that we encountered a media-specific artifact from ½X MS. It is possible that ROS or O_2_ could inhibit nitrogen assimilation enzymes. Alternatively, during standing incubation *B. subtilis* enters a biofilm lifestyle that may alter inorganic nitrogen assimilation (70), whereas agitation introduces shear stress and limits biofilm formation. Finally, another possibility is that the energetic or redox costs of building biomass from inorganic nitrogen are simply too high for an individual vegetative cell and that either provision of an organic nitrogen source or formation of a biofilm are needed to overcome this cost. Looking ahead, further interrogation of each of these possible contributions should help design hydroponic processes that can better support growth of engineered *B. subtilis* bioinoculants, and our findings may also guide media design when using *B. subtilis* during aerobic fermentation.

## Methods

### Materials and Reagents

The following was purchased from MilliporeSigma (Darmstadt, DL): Magnesium sulfate, Murashige Skoog (MS) basal media, KOD Xtreme Hot Start DNA Polymerase, trifluoroacetic acid, acetonitrile, monopotassium phosphate, dipotassium phosphate, magnesium sulfate, calcium chloride, Tris base, chloramphenicol, ammonium sulfate, magnesium chloride, thiamine-HCl, EDTA, NAD, Cobalt (II) Chloride hexahydrate, sodium molybdate dihydrate, L-valine, L-isoleucine, L-leucine, Potassium L-glutamate, casein hydrolysate.

The following was purchased from Thermo Fisher Scientific: potassium nitrate, trisodium citrate dihydrate, manganese chloride, zinc chloride, copper chloride dihydrate, ammonium iron citrate, glucose, sucrose, Iron (III) Chloride, LB agar, ferric ammonium citrate, agarose, sodium hydroxide, HPLC grade water, SybrSafe.

Phusion polymerase, T5 exonuclease, Taq DNA Ligase, and SSB were purchased from New England Biolabs (Ipswich, MA, USA). Hydrochloric acid was purchased from RICCA (Arlington, TX, USA). LB was purchased from RPI (Mount Prospect, IL, USA). L-aspartate, ammonium chloride and MOPS were purchased from TCI. From Integrated DNA Technologies (IDT) (Coralville, IA, USA), primers were purchased. Surfactin was purchased from Smolecule (San Antonio, TX, USA).

### Media Preparation

MC media was made from a 10x stock concentration, which has final concentrations of 1M potassium phosphate pH 7, 30 mM sodium citrate, 20% (w/w) glucose, 220 mg/mL ferric ammonium citrate, 1% casein hydrolysate, 2% potassium glutamate. Aliquots were stored at - 80°C. MC media was made for transformations and supplemented with 3mM MgSO_4_. TE buffer for colony PCRs was composed of 10 mM Tris-HCl and 1 mM EDTA, pH 8.0. LB and LB agar had final concentrations of 10 g/L casein digest peptone, 5 g/L yeast extract, 10 g/L sodium chloride. LB agar also had an additional 15 g/L agar. Media was then sterilized via autoclave.

S750 media was made in a total volume of 50 mL, by mixing 5 mL 10X S750 salts (recipe below), 0.5 mL 100× S750 metals (recipe below), 0.5 mL 1 M glutamate and 2.5 mL 20% glucose, all in ddH_2_O and filter sterilized (not autoclaved). 10X S750 salts were made in 100 mL aliquots. 10.47 g MOPS (Free acid), 1.32 g ammonium sulfate (NH_4_)_2_SO_4_, 0.68 g potassium phosphate monobasic and buffered to pH=7 with 50% potassium hydroxide. Media was filter sterilized and stored at 4 °C. 100x S750 metals have final concentrations of 0.2 M MgCl_2_, 70 mM CaCl_2_, 5 mM MnCl_2_, 0.1 mM ZnCl_2_, 100 microgram/mL thiamine-HCl, 2 mM HCl, 0.5 mM FeCl_3_. Iron was added last to prevent precipitation. Media was sterilized and stored in black falcon tubes 4 °C.

Spizizen minimal media contained 6 g/L monopotassium phosphate, 14 g/L dipotassium phosphate, 1.25 g/L trisodium citrate dihydrate, and 0.2g/L MgSO_4_·7H_2_O. Trace elements were supplemented from 100x stock containing 0.55 g/L CaCl_2_, 0.1 g/L MnCl_2_·4H_2_O, 0.17 g/L ZnCl_2_, 0.033g/L CuCl_2_·2H_2_O, 0.06 g/L CoCl_2_·6H_2_O, and 0.06 g/L Na_2_MoO_4_·2H_2_O. As well as 100x F-Citrate Solution containing 0.0135 g FeCl_3_·6H_2_O and 0.1g trisodium citrate dihydrate. Sucrose was supplemented at a final concentration of 1.5%. Nitrogen was added depending on experimental condition, but importantly no casein hydrolysate was added to the base media composition. Supplemented nitrogen sources (ammonium chloride, potassium nitrate, L-glutamate, L-glutamine, L-valine, L-leucine, L-isoleucine, and L-aspartate) were made at in 1 M stock solutions and filter sterilized through 0.22 µm syringe filter before adding to media. Branched chain amino acids were titrated with NaOH until soluble before filter sterilizing.

### Strain Generation

Polymerase chain reaction (PCR) based DNA amplification was performed with KOD Xtreme Hot Start Polymerase. PCR primers were designed to have 20 bp of overlap for ligation via Gibson isothermal assembly. Isothermal reactions were done in 13 µL final volume homemade Gibson mix that was prepared according to the original 2009 Gibson paper(71), at 50°C for 1 h. The assembled construct was PCR amplified before purification and then used for transformation. The gene encoding *sfp* was purchased from the Bacillus Genus Stock Center (BGSCID: ECE221). Primers used in this study can be found in **Table S1,** as well as additional information on strain construction.

Transformation was done by inoculating 1 mL of modified competence (MC) media from plates. At 4 h, the DNA was added to 200 µL cells and grown at 37°C for 2 more hours before plating on selective media. Colonies were verified to have the DNA by colony PCR, where cells were suspended in 50 μL TE buffer +10 μm glass beads, then vortexed for 10 min, boiled for 30 min, vortexed for 10 min. 1 μL was used as a PCR template. Purified colony PCR DNA was sequenced by Plasmidsaurus to confirm genotypes. A list of strains and plasmids used can be found in **Table S2**.

PY79 sfp^+^ was constructed by amplifying 2 pieces from bDS176 (72) with primers: oAC0001 and oMAJ0282, and oAC0015 and oAC0008. The first PCR product contained the upstream homology, antibiotic resistance marker, and P_veg_ promoter and the second contained the downstream homology. Sfp was amplified from pSV-sfp using primers oAC0014 and oAC0005. Pieces were assembled via Gibson assembly and amplified with oAC0001 and oAC0008. Purified PCR was used to transform PY79.

### Strain Growth Assays

Strains were grown from glycerol stocks in LB to an OD of approximately 1.0 before washing 5x via centrifugation in PBS. The OD was measured after washing with a spectrophotometer at a wavelength of 600 nm. Strains were then inoculated in triplicate in the tested media with at an initial OD of 0.1. Timepoints were grown in 96 well plates, with round bottom for shaken conditions and flat bottom for static conditions. 200 µL of sample was moved to a clear, flat bottom plate before measuring OD_600_ on a Spectramax i3x. Continuous growth assays were performed in clear bottom, flat 96-well plates. OD_600_ was measured every 10 min with medium plate shaking between reads over 24 hours, grown at 37°C.

### Growth and Surfactin Production Assays

Strains were grown from glycerol stocks in LB to an OD_600_ of approximately 1.0 before washing 5x via centrifugation in PBS. The OD was measured after washing with a spectrophotometer at a wavelength of 600 nm. Strains were then inoculated in triplicate for each of the relevant media conditions at the relevant OD_600_, 0.01 for low inoculation OD, 0.1 for growth assays, or 0.2 for high inoculation OD. Media conditions varied by media (LB, S750, ½X MS, and Spizizen media), where nitrogen sources were varied in ½X MS and Spizizen media to test the influence of organic or inorganic nitrogen sources. Strains were grown with appropriate antibiotics, 5 μg/mL chloramphenicol for PY79 sfp^+^ or none for wild-type strains. Strains were grown in an incubator to maintain the relevant temperature. Shaking conditions were done at 1000 rpm in 96 well plates or 250 rpm for culture tubes, strains grown under static conditions were grown in incubators with no shaking. *B. subtilis*168 was supplied with 50 μg/mL tryptophan when grown in minimal media. Timepoint OD_600_ values were measured in a Spectramax i3x plate reader at 600 nm, where 200 μL of culture was moved to a clear, flat bottom 96 well plate. For surfactin analysis, timepoint OD_600_ was measured before 200 μL of culture was centrifuged to pellet cells and supernatant was collected for HPLC analysis.

Continuous growth assays were performed using a Spectramax i3x. Cells were innoculated at an initial OD_600_ of 0.1 in clear bottom, flat 96-well plates. OD_600_ was measured every 10 min with medium plate shaking between reads over 24 hours, grown at 37°C.

### Surfactin Quantification

Surfactin quantification was done via high performance liquid chromatography with an Agilent 1100 Infinity model with a Zorbax Eclipse Plus-C18 column with a guard column installed. To isolate surfactin peaks, an isocratic method of acetonitrile with 1% trifluoroacetic acid was used. 50 μL of sample was injected and run for 10 minutes, and absorbance was analyzed at 205 nm. Quantification was accomplished by comparing peak areas of three major peaks in the commercial standard.

### Transcriptome Analysis

Transcriptomics were performed on data from Nicholas et al, comparing conditions of BT, B36, and B60, considering genes relevant to nitrogen assimilation (57, 73–78). Genes associated with nitrogen transport, regulation, and metabolism were analyzed. Log_2_ fold change was calculated by the ratio of log base 2 of the mean transcript level reported in the B36 condition versus the BT condition reported in Nicolas et al (56).

## Acknowledgements

AMK and HPB acknowledge support from the U.S. Department of Agriculture (USDA) National Institute of Food and Agriculture (NIFA) Biotechnology Risk Assessment Grants (BRAG) program, Award No. 2021-33522-35807. AMK acknowledges support from the Foundation for Food and Agriculture Research (FFAR) under their New Innovator Award, which is Award No. FF-NIA21-0000000060. RW acknowledges support from the Microbiology Graduate Program at University of Delaware. W23 strain used in this work was provided by the USDA-ARS Culture Collection (NRRL).

